# Novel Pore Size-Controlled, Susceptibility Matched, 3D-Printed MRI Phantoms

**DOI:** 10.1101/2022.10.10.511634

**Authors:** Velencia J. Witherspoon, Michal E Komlosh, Dan Benjamini, Evren Özarslan, Nickolay Lavrik, Peter J. Basser

## Abstract

Diffusion magnetic resonance imaging (dMRI) methods are commonly employed to infer changes in tissue microstructure. Quantities like the apparent diffusion coefficient (mADC), and the fractional anisotropy (FA), derived from diffusion tensor imaging (DTI) data, characterize voxel-averaged diffusion properties, whereas double pulse field gradient (dPFG) or double diffusion encoded (DDE) MR methods can be used to characterize heterogeneous diffusion processes occurring within the voxel. Owing to its unique modular design, our novel 3D-printed dMRI phantom exhibits both macroscopic and microscopic anisotropy and can serve to calibrate measures of them. Our phantom susceptibility is close to that of water’s, enabling fast diffusion weighted echo-planar image (DW-EPI) acquisitions to be used to scan it. 3D printed microstructures offer a new medium with which to vet and validate theoretical models of diffusion and pipelines used to estimate it.

**Highlights:**

- Research highlight 1: We report the design concept and fabrication of dimensionally stable, uniformly oriented blocks or modules that can be assembled into large-scale MRI phantoms. Waffle-like structures containing blocks of aligned microcapillaries can be stacked into even larger arrays to construct diameter distribution phantoms, or fractured, to create a “powder-averaged” emulsion of randomly oriented blocks.
- Research highlight 2: This phantom can be used to vet and calibrate various MRI methods, such as DTI, AxCaliber MRI, MAP-MRI, and various multiple pulsed field gradient (PFG) or multiple diffusion-encoded microstructure imaging methods.

**Graphical Abstract:** 

## 1. Introduction

Microstructure imaging methods are commonly employed in a variety of biological and clinical applications. In the former case, these include elucidating features of brain structure and organization, which are often used to determine anatomical connectivity, for instance in neuroscience. In the latter case, these include assessing normal and abnormal developmental processes, pathological tissue states, or following features of normal and abnormal aging or detection of trauma, *inter alia*. A goal of microstructure imaging is to use macroscopic scale diffusion-weighted imaging (DWI), and potentially other imaging data, in conjunction with physical models of the underlying diffusion process at finer length scales to infer key microstructural features and morphological parameters, often in brain white matter, such as mean axon diameter (e.g., in dPFG MRI [1, 2] or the axon diameter distribution, e.g., in AxCaliber MRI [3], DKI [4], MAP-MRI [5]. There is a critical need to design and create MRI phantoms that can be used to test and calibrate these microstructural imaging methods and models and vet their associated data processing and display pipelines. Generally, MR phantoms play a crucial role to test data acquisition and analysis pipelines, ensuring that model results conform to experimental data. In clinical and pre-clinical scanners, MRI phantoms can be used to help ensure precision and accuracy of MRI data in single subjects, as well as in longitudinal scans and in multi-site studies [6]. One way to vet such pipelines is to see whether they recapitulate the “ground truth” when they are subjected to data obtained from structures whose morphological features such as pore size and shape, connectivity, etc., are known with certainty and can tax the estimate of model parameters. As diffusion-related parameters are increasingly applied to assess more complex tissue morphologies, it is vital to assess the performance of these methods on model phantom/material systems that mimic salient features of tissue, but nonetheless are simple enough that analytical or approximate solutions can be found to describe the signal attenuation profiles measured within these structures.

In attempting to infer features of gray matter, in which there are many crossing processes and fibers oriented somewhat randomly, there is a need to develop MR techniques that can interrogate these powder-averaged microscopically anisotropic structures. Models and experimental pipelines used to estimate pore size distributions in such complex media require the development of phantoms or ground truth structures that can tax such methods [7], Since Cheng and Cory[8] first described and measured microscopic anisotropy within ellipsoidal yeast cells that were randomly oriented in an emulsion, numerous studies have investigated the relationship between the mPFG diffusion signal attenuation profile on pulse sequence parameters and microstructural features, such as distributions of compartment shape, size, and orientation (Ozarslan, 2009[9]). The development by Grebenkov of the multiple correlation functions framework [9, 10] has led to the study of simple pore geometries and arbitrary orientations from NMR diffusion signals, particularly to tease out features of microscopic anisotropy in powder-averaged structures. In attempting to transition to microstructure MRI in tissue, particularly in *in vivo*, we must be able to calibrate MRI scanners and standardize instrument performance. The use of designer phantoms greatly aids in this process. Several MRI phantoms have been designed and fabricated [11, 12, 13, 14, 15, 16, 17, 18] to represent microscopically anisotropic but macroscopically isotropic grey matter tissue. These phantoms, although functional, often suffer from susceptibility contrast between the phantom material and water as compared to biological tissues, safety concerns, such as spillage of organic solvents, or fragility, limiting phantom transportability and robustness. To address these, we have shown that advanced 3D printing technology with high-accuracy and precision fabrication techniques can be utilised to produce MRI phantoms that are anisotropic, can be randomly oriented, and have magnetic susceptibilities close to water, thus, providing model systems mimicking aspects of the anisotropic microcapillary-like structures found in gray and white matter in the central nervous system (CNS).

## 2. Experimental Methods

### 2.1. 3D Printed Phantom

MRI Phantoms were printed using a Photonic Professional GT stereolithography tool based on 2-photon polymerization (Nanoscribe, GmbH) at the Center for Nanophase Materials Sciences (CNMS) at Oak Ridge National Laboratory (ORNL). This printer was chosen because of its high resolution and ability to print features with high fidelity (+*/−*100*nm*). The 3D computer assisted designs (CAD) of capillary arrays arranged in hexagonal packing within 300*µm* x 300*µm* x 300*µm* cube/blocks were compiled using the 3D graphic editor built into COMSOL Multiphysics software (Figure 1). Hexagonal packing and the smallest feasible inter-capillary distances were used to maximize the pore volume and thus the signal-to-noise ratio (SNR) of the MRI. Four small parallelograms were added to the bottom side of each block to elevate capillary arrays and facilitate removal of un-crosslinked photo-resin from the high-aspect-ratio capillaries during the development step. The compiled 3D CAD of arrays of capillaries was saved in the STL format and subsequently imported into DeScribe software (Nanoscribe GmbH) to compile job files with various printing parameters tabulated in 2.

**Figure 1:**
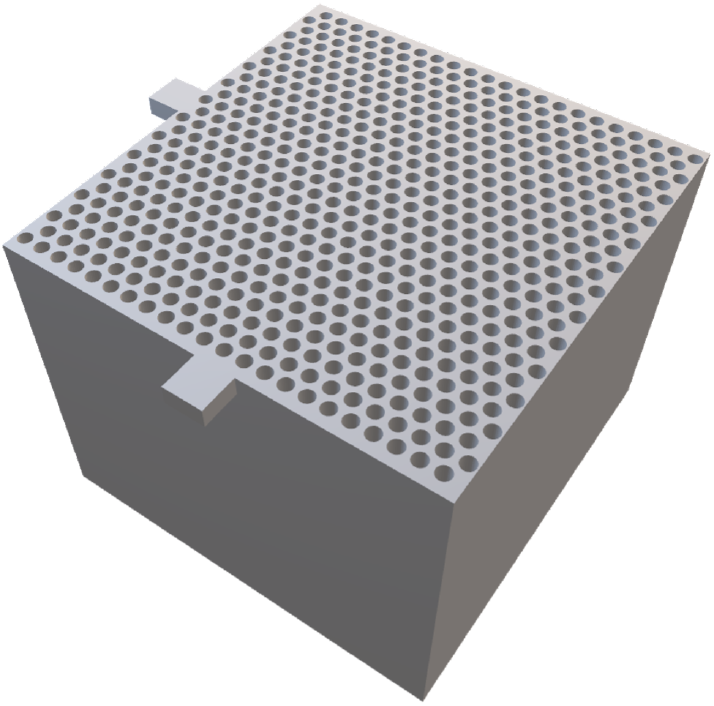
CAD schematic of printed blocks produced using COMSOL Multiphysics software.

The slicing and hatching printing parameters were varied and optimized in order to produce blocks with solid and smooth walls between adjacent capillaries. To meet the requirement of the targeted 10*µm* inner diameter (ID), the diameters of capillaries in final CAD were adjusted and after several iterations of printing tests. IP-S photo-resin, a poly (methyl methacrylate) based formula, (Nanoscribe, GmbH) was used for all printing tasks in this study. This printer was chosen because of its high resolution and small tolerance (+/- 100nm). The finalized design was intended to meet the above criteria for capillary arrays of 10*µ*m inner diameter(ID), contained within cube/blocks of 300*µ*m x 300*µ*m x 300*µ*m.

To accumulate total capillary volumes sufficient MRI characterization, job files contained step-and-repeat loops compiled in such way that each sample contained 7×7 to 15×15 2D arrays of individual blocks shown in Figure 1. 150*µ*m thick circular cover slips or 500*µ*m thick silicon chips were used as substrates for printing. Development of the printed structures was carried out in 1-methoxy-2-propyl acetate for at least 24 hours with subsequent rinsing in isopropanol and drying in a gentle stream of filtered dry nitrogen. A 24-48 hour development time was found to be necessary to ensure complete removal of the uncrosslinked IP-S photoresin from the high-aspect-ratio capillaries.

### 2.2. Microscopy Characterization and Sample Preparation

Scanning electron microscopy (SEM) images were taken before and after the MRI experiments on a Hitachi instrument. The blocks were coated with a 5-10nm layer of gold when imaged after the MRI acquisition. The images were then analyzed with ImageJ to determine the microscopic pore size distribution. The blocks were plasma treated to make the surface hydrophilic and then loaded with water in a glass tube. The tube was place under vacuum for 5-7 days to eliminate air bubbles from the pores and surfaces which would create large susceptibility gradients if not removed, and the sample is then sealed for imaging. Susceptibility-matched plugs were placed at the bottom of the glass tube and on top of the sample. This also prevented movement of the block during the duration of the experiment, while suppressing susceptibility gradients from being generated at the top and bottom surfaces of the phantom, as well.

### 2.3. MRI Measurements and Analysis

A mixed model of restricted and Gaussian diffusion [3] was used to fit the dPFG data, with the restricted diffusion modeled to arise from a cylindrical pore geometry [25]. We utilized custom MATLAB code to estimate the voxel-wise average pore size accounting for the diffusion gradients profile.

The 3D printed capillary blocks were then removed from their support, randomly distributed in a glass tube, loaded with H_2_O, and re-measured using a diffusion tensor imaging (DTI) [26] protocol with a diffusion-weighted 3D-EPI acquisition. Inversion Recovery preparation for FLASH3D experiments[27] was used to produce the high resolution *T*_1_-MAP, while spin echo Fast low angle shot (FLASH3D) and multi-slice multi-echo sequence (MSME) were utilized to produce *T*_2_-MAPs. The measured relaxation time constants were used to set and constrain the diffusion time and echo time. Additionally, a diffusion weighted *T*_1_ Map and diffusion-weighted *T*_2_ were measured with an inversion recovery preparation sequence and an varying the echo time of the EPI acquisition. These were acquired to isolate the relaxation behavior of the protons signals that would contribute to the d-PFG and DTI experiments. These results are located in the SI. The experimental parameters can be found in 3.

### 2.4. Double Pulse Field Gradient MRI

Before creating the randomly oriented phantom, the aligned capillary pore size was measured using a double-pulsed field gradient (dPFG), also known as a double diffusion encoding (DDE) spin echo imaging sequence, using a previously proposed protocol [28]. The aligned MRI data were acquired with dPFG NMR followed by a 2D spin echo MRI sequence with the following parameters where the axial slice thickness = 200 *µm*, and spatial resolution = 50 × 50 × 200 *µm*^3^, 2D dPFG NMR parameters were as follows: *TE* = 35 ms, *τ*_*m*_ = 0; *δ* = 3 ms; Δ = 30 ms; *ϕ*, the angle between the two PFG blocks, varied from 0 to 360 with intervals of 15 degrees in the restricted plane, with q range from 0 to 5.73 *x*10^4^ *m*^−1^. These same blocks were then removed from their support, randomly suspended in 4 mm a glass tube, and re-scanned using an adjusted dPFG NMR protocol [29] and diffusion tensor imaging (DTI) [26]. The randomly oriented phantom data were acquired dPFG parameters were as follows *tau*_*m*_ = 0 or 6 ms; *δ* = 3 ms; Δ = 30 ms; Three shells of 7, 16, and 37 directions with q ranging from 0 to 8.5947 *x*10^4^ *m*^−1^. For the *tau*_*m*_ = 0 experiment the angles describing the gradient directions and the shell number were varied using a previously published 3D acquisition scheme[30]. For the long *tau*_*m*_ acquisition each direction was applied with *ϕ* = 0 and 180 degrees.

### 2.5. Additional MRI Considerations

The randomly oriented wafer phantom was scanned using a standard Point RESolved Spectroscopy (PRESS) sequence where the volume for individual experiments were registered to different block orientations and free water region of interest identified via DTI. We used TORTOISE [31, 32] to produce Diffusion-Encoded Color (DEC) and Fractional Anisotropy (FA) images from the DTI experiments. The *T*_1_MAP and *T*_2_MAP were produced and registered to the DEC image since the same resolution and field-of-view (FOV) were utilized for all experiments. Using a customized MATLAB code the voxel-wise decays were fitted to a mono-exponential with a constant baseline using a non-linear least-squares fitting routine to identify the best fit time constants. To qualify how the phantom material’s susceptibility may affect our diffusion imaging sequence, we conducted multiple EPI experiments varying the number of segments and quantitatively compared the images to one another. Experimental details and parameters can be found in 3. All MRI measurements were performed on a 7T Bruker MRI scanner with an AvanceIII spectrometer and a Micro2.5 microimaging probe with a 5 mm ^1^H RF coil.

## 3. Results

The tested printing speeds and the resulting wall uniformity are shown in Table 2 and figure 2. The finalized MRI Phantom printed under the conditions listed in 2 for Sample C in Table 2. This print resulted in an array of aligned blocks, see 3 where the gaps between adjacent block are approximately 50 *µm*. This aligned phantom was investigated by dPFG and signal from the 200 *µm* slice was fitted to a mixed Gaussian model to calculate the size of the restriction. The resulting voxelwise pore size estimate from the aligned dPFG resulted in fairly homogeneous pore size values, with mean and standard deviation of 8.38 +*/−* 1.45 *µ*m (7), While the nominal value of 10.7 +*/−* 0.323 *µ*m was obtained from pore size analysis of several SEM images using ImageJ. The blocks were dispersed in water in a 10 mm NMR tube and investigated via *T*_1_ weighted FLASH 3D experiments where the inversion time was varied. The *T*_1_ weighted image, Panel B, was acquired with a 13 ms recovery time **??** where Panel A in figure **??** results from a series of FLASH3D experiments with varied inversion recovery preparation(s) that were compiled and analyzed to generate a T_1_ map with custom MATLAB code. Figure 4 is the result of a high-resolution segmented EPI acquisition with varying k-space segments: 2, 4, and 8. The 4 and 8 segment images were compared to the 2 segment image using the structural similarity index measure (SSIM) index and resulted in high SSIM values of 0.9984 and 0.9661 respectively. Figure **??** is the result of a series of FLASH3D experiments with a 13 ms inversion recovery preparation. In Figure 6, Axial slice showing high resolution (50×50×50 *µm*) DEC map and the FA map using DTI data. This shows that the blocks are randomly oriented and microscopically anisotropic while the SEM in Figure 5 shows that they are microscopically anisotropic with FA well above 0.75.In Figure 6, Diffusion-Encoded-Color (DEC) map (* cite pajevic and pierpaoli*) using DTI data, shows that the blocks are randomly oriented but microscopically anisotropic. The SEM in Figure 3 also show that they are microscopically anisotropic. Figure (to be inserted) shows blocks (A-G) from the center slice chosen at random where Point RESolved Spectroscopy (PRESS) [34]) was performed to record the chemical shift behavior as a function of orientation.

**Table 1:** Previous Microscopic Anisotropy Phantoms

| Sample Description | Material | Ref | Size $\mu m$ | Shape | Robustness |
| --- | --- | --- | --- | --- | --- |
| Water-filled hollow tubes | Fused Silica Fiber | [19, 20] | I.D.: 9, 20, O.D.:150 | Cylinders | Low SNR, susceptibility challenges |
| Silicon oil-filled capillary bundles | Glass | [13] | 5,10,25 | Cylinder Array | Transportable, stable, susceptibility challenges |
| Water Saturated Wood Phantoms | Biological | [18] | Varied | Lamellar Layers | Variables in wood species, not sustainable |
| Liquid Crystal | water, detergent, and hydrocarbon | [14] | 0.007 | Cylinder Array | Non-relevant sizing |
| Capillaries | Glass | [21] | 23, 48, 82 | cylinders | Susceptibility Challenged |
| Hollow Fibers | Polypropylene | [22] | I.D. 12 O.D. 33.5 | Hollow cylinders | Porous wall with a large non-anisotropic restricted diffusion population |
| Plants | Asparagus, Celery | [23, 24] | 100 | cylinder and cells | not stable or repeatable from sample to sample |

**Table 2:**
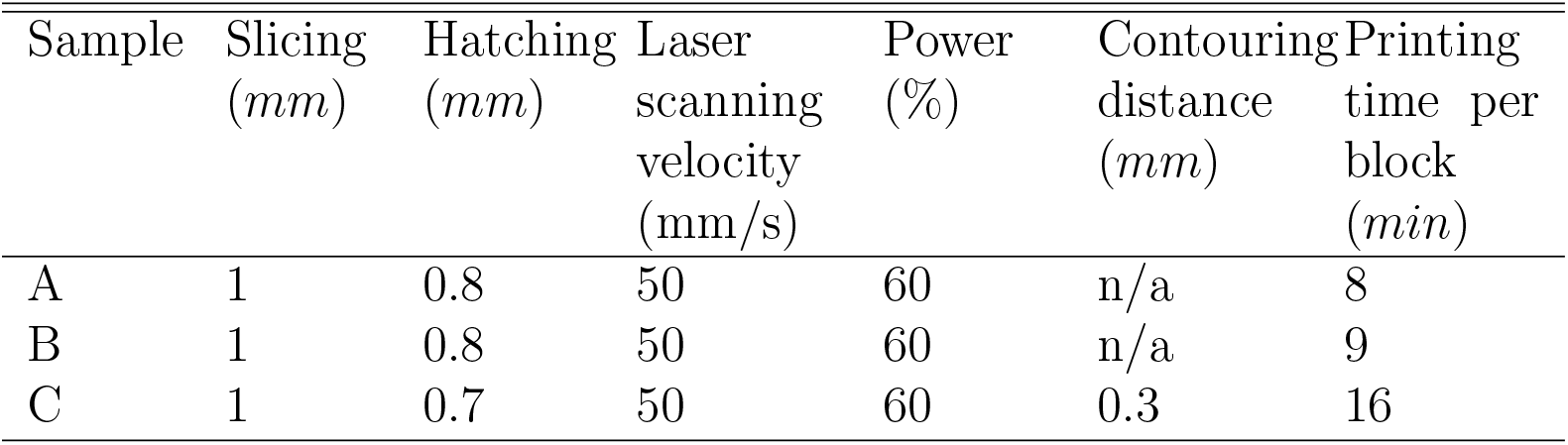
Printing Conditions

**Table 3:**
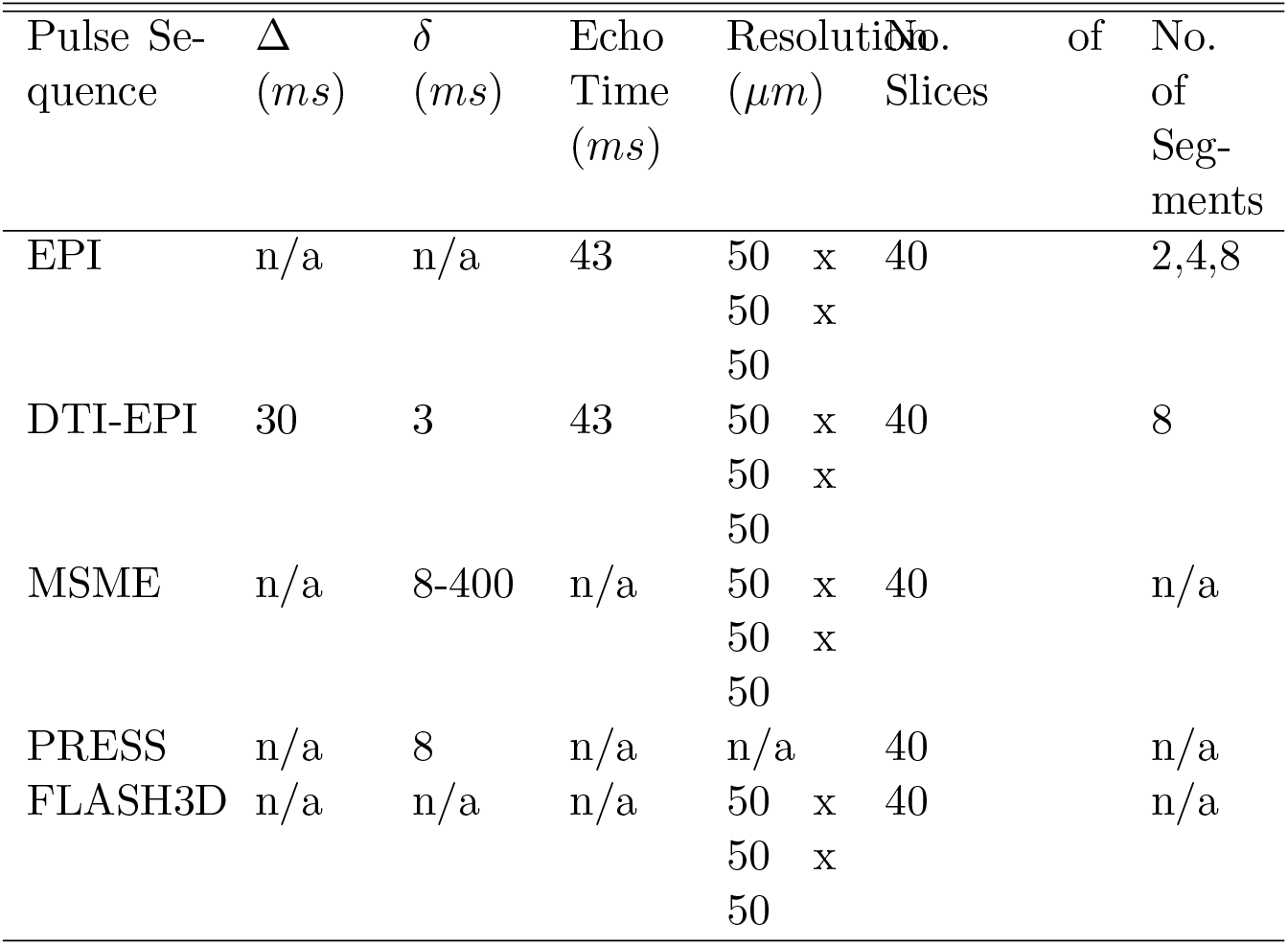
Experimental parameter values

**Table 4:**
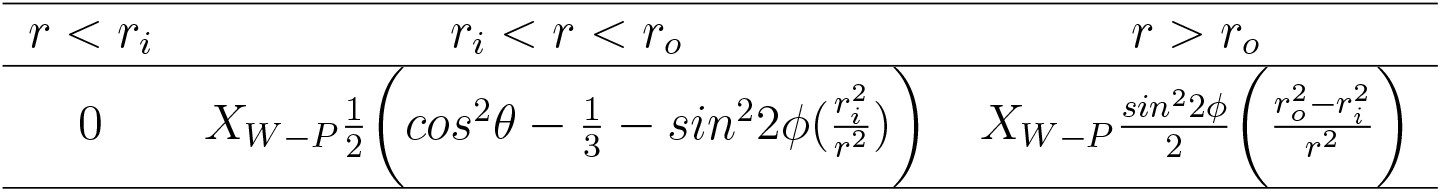
Frequency perturbation in hollow cylinder for isotropic susceptibility. Abbreviation *X*_*W−P*_ for *X*_*H*2*O*_ *− X*_*P MMA*_ used below

| $r < r_i$ | $r_i < r < r_o$ | $r > r_o$ |
| --- | --- | --- |
| 0 | $X_{W-P} \frac{1}{2} \left( \cos^2 \theta - \frac{1}{3} - \sin^2 2\phi \left( \frac{r_i^2}{r^2} \right) \right)$ | $X_{W-P} \frac{\sin^2 2\phi}{2} \left( \frac{r_o^2 - r_i^2}{r^2} \right)$ |

**Table 5:** Model Parameter values

| Model Fit | $D_{spp}$ | Cylinder Diameter $\mu m$ | Free Population | Bound Population |
| --- | --- | --- | --- | --- |
| Free + Restrictions | 1.85 | $7.80 \pm 0.08$ | $0.60 \pm 0.005$ | – |
| Free + Restriction + Bound | 1.85 | $8.22 \pm 0.06$ | $0.60 \pm 0.005$ | $0.0037 \pm 0.0007$ |

**Figure 2:**
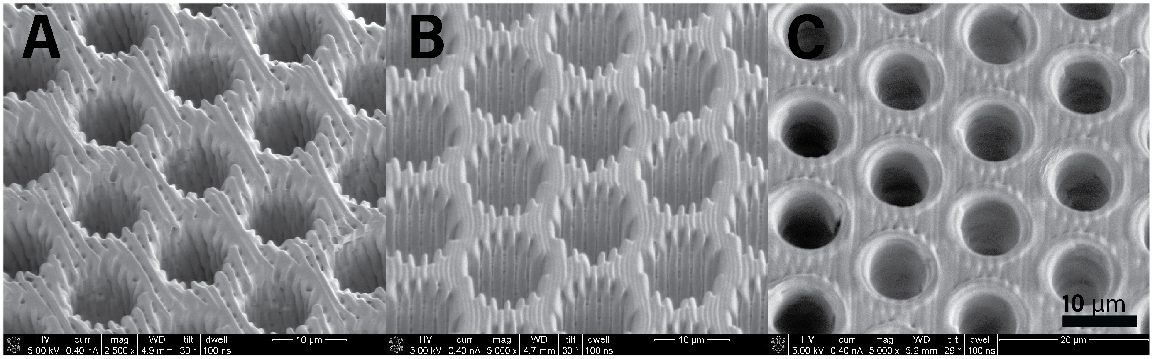
SEM images acquired at CNRM of singular block printed at the condition listed in tab:printcond for samples A, B, C

**Figure 3:**
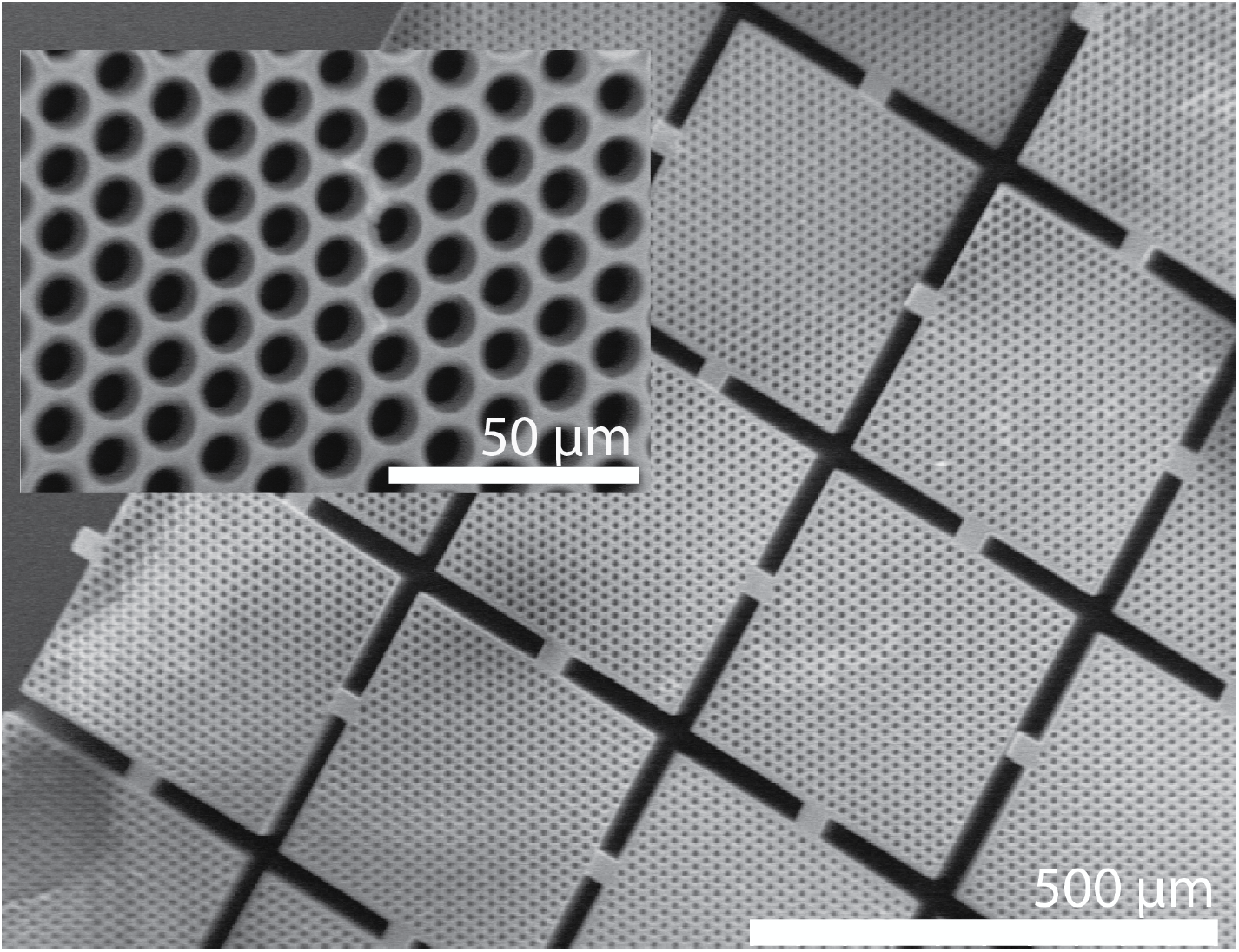
SEM images acquire of finalized MRI Phantom NIH with the white line indicating the scale bar for 500 *µm* and the inset scaled bar for 50 *µm*

**Figure 4:**
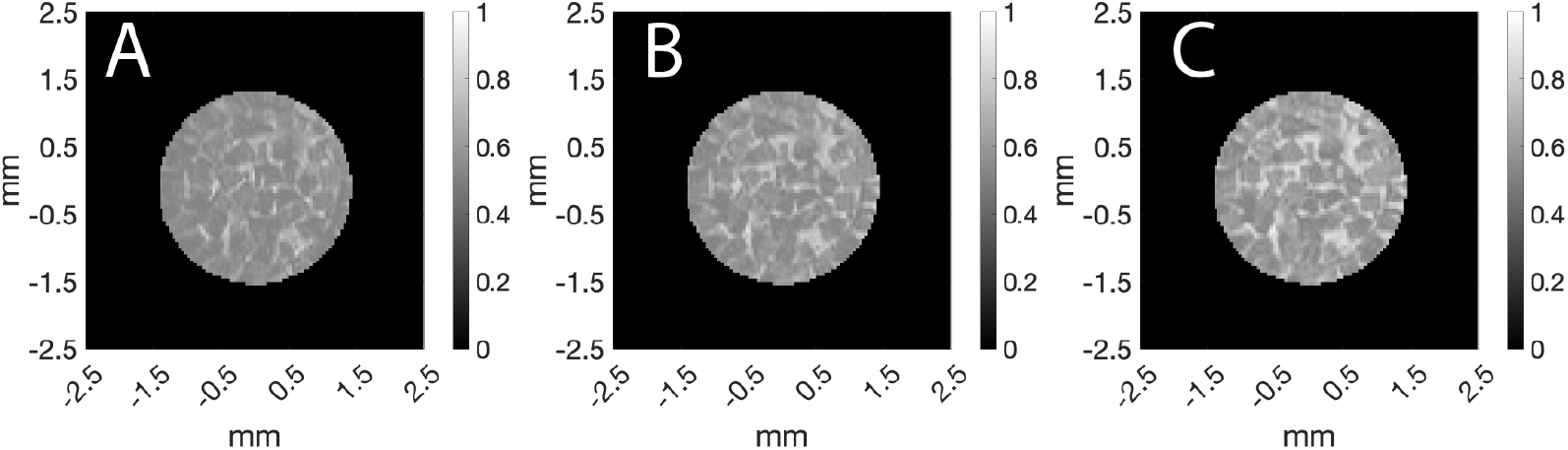
The same axial slices acquired using an EPI sequence where the number of segments were varied from 2 (A),4(B), and 6(C) segments, with TE = 54 ms. the 3D voxel size is 50 *µ*m^3^. The relative SSIMs of images B and C to the first image A are 0.9984 and 0.9661, respectively.

**Figure 5:**
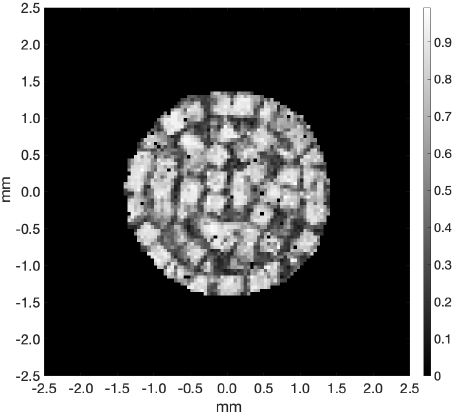
Axial slice of the FA map calculated using TORTOISE, with 50*×* 50 *×*50 *µ*m^3^voxel size, demonstrating that diffusion in each block is highly anisotropic with FA *>* 0.75.

**Figure 6:**
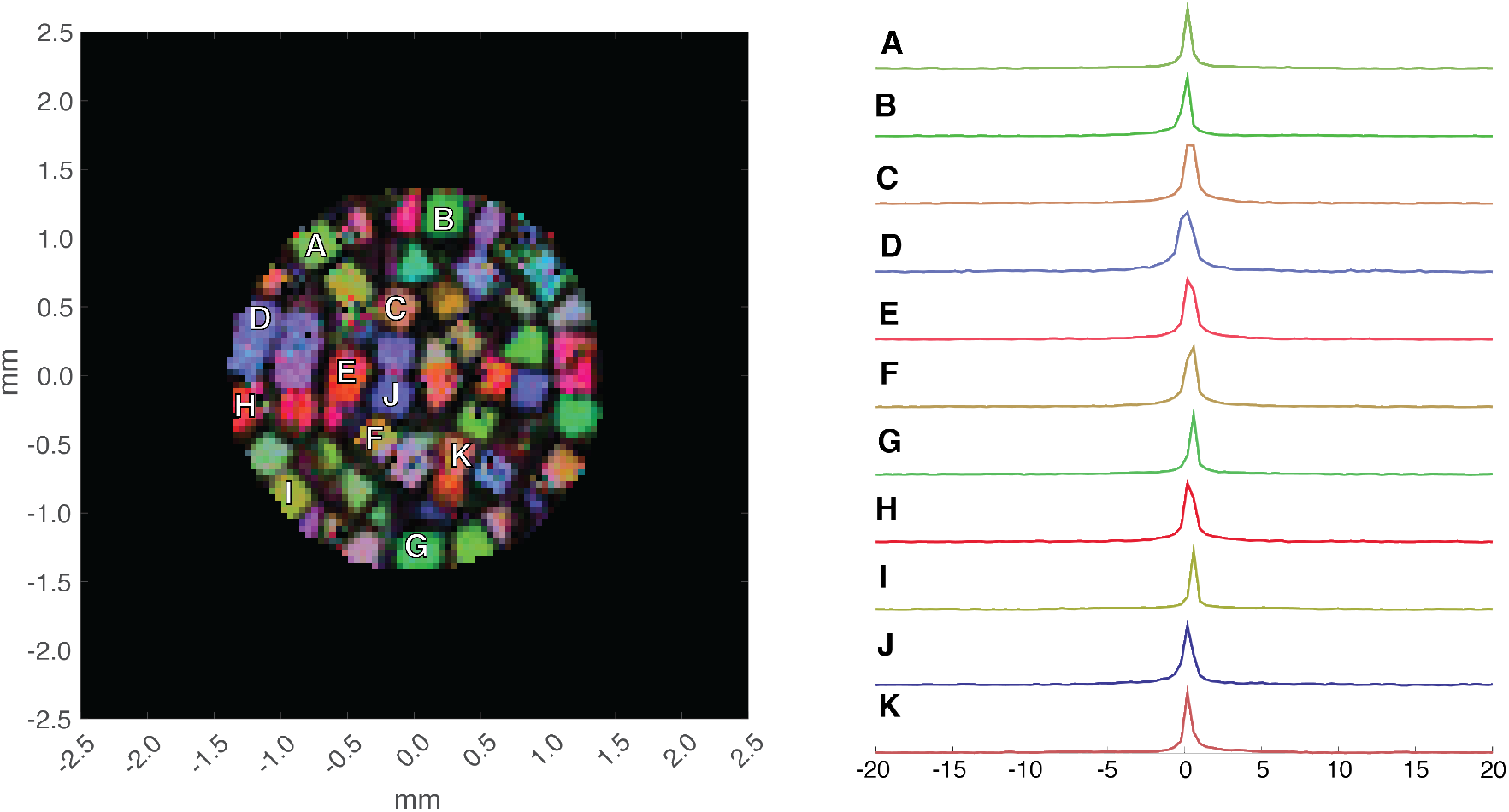
The left panel is the Diffusion-Encoded Color (DEC) MRI map from a TORTOISE analysis obtained with 50 *µm*^3^ isotropic voxels. The letters on the left image represent voxels containing blocks with different orientations, for which PRESS ^1^H spectra are also obtained. The corresponding spectra are shown on the right panel plotted against *ω*_0_, the Larmor frequency (in ppm) and are virtually indistinguishable within the instrument resolution. If there were significant susceptibility artifacts, one would expect to see the Larmor frequency vary with orientation of the capillaries with respect to the main magnetic field direction.

**Figure 7:**
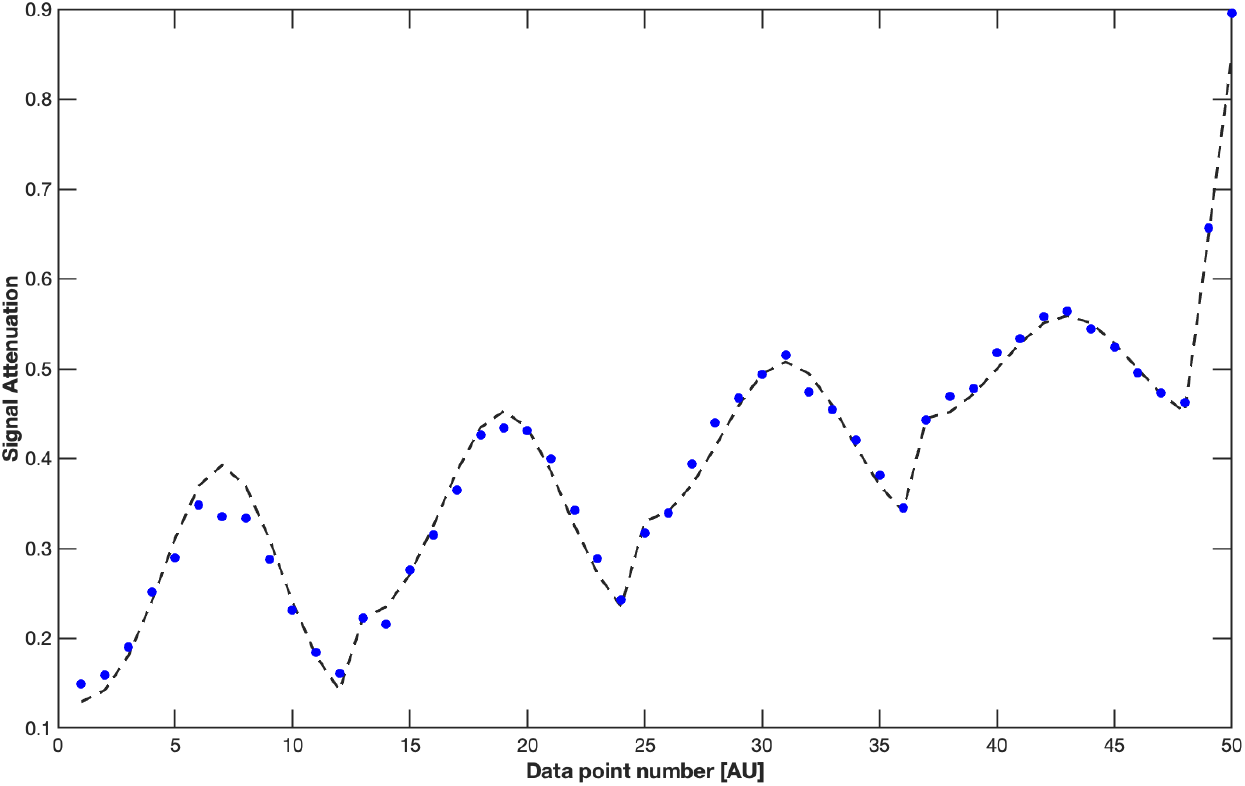
The experimental signal intensity and the resulting fit (Mixed Gaussian and restriction model) [33] are plotted against arbitrary parameters of varying applied pulse gradient strengths and directions. The resulting diameter for the aligned blocks, diameter = 8.58 *µm*, with an apparent diffusivity of 2.0 × 10^*−*9^ m^2^/s, and the fraction of the freewater signal associated with the Gaussian distribution, *f*_*g*_ = 0.38.

**Figure 8:**
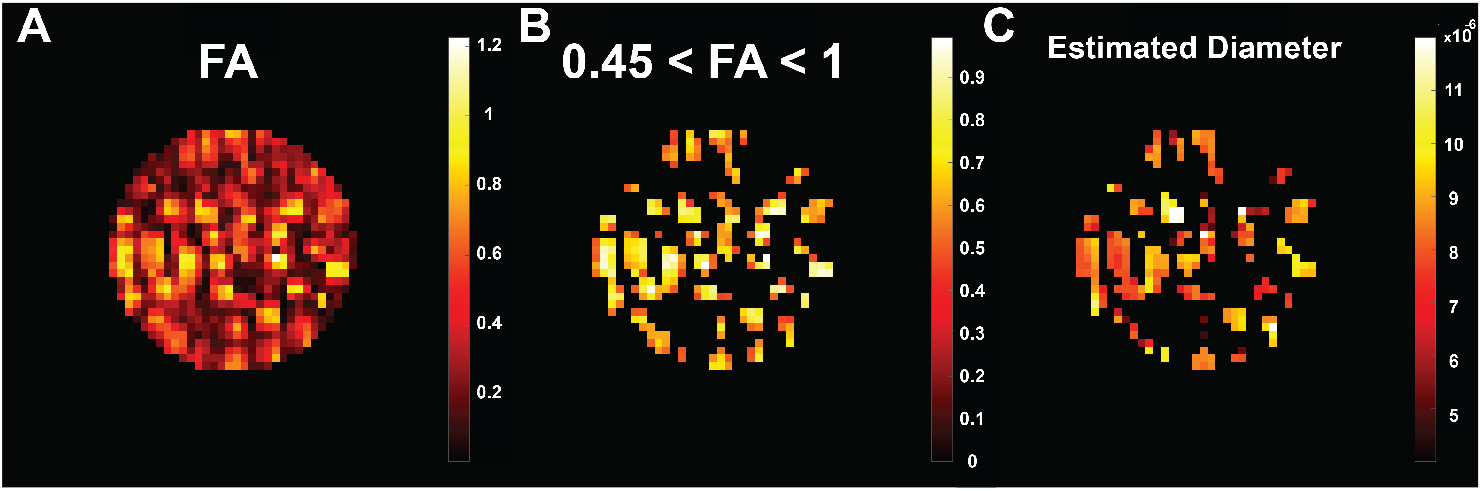
100*µ* isotropic resolution images, showing (A) FA, (B) FA with an applied threshold of FA*>*.45, and (C) the calculated diameter map using the framework from Benjamini et al. [28]

## 4. Discussion

### 4.1. 3D Printing Design

At the outset, we sought to satisfy basic criteria required for a microscopically anisotropic but macroscopically isotropic phantom. We needed to print a microscopic structure that has high morphological uniformity, and exhibits restricted diffusion that would result in high diffusion anisotropy (FA *>* 0.6), and would have a magnetic susceptibility close to H_2_O. This would ensure that the voxels are easily distinguishable from free water in a basic DTI acquisition. It was desirable for the pore fraction to be as close to its maximum value (72 %) as possible to ensure that the signal to noise (SNR) is sufficiently high so that different models could robustly be fit to experimental data. The phantom substrate needed to have a magnetic susceptibility similar to that of water to enable use of diffusion weighted 3D EPI and fast acquisition schemes commonly used in clinical settings and at high magnetic field without any imaging or other artifacts. There should be no exchange of water between pores to simplify the model that could be used for validation. Such phantoms should also be stable, and withstand degradation or change to the NMR observables for at least a span of 2 years. 10 microns, close to the detection limit for clinical MRI scanners, was set as the target inner diameter for the phantom’s pores. The pitch of the capillaries was chosen to ensure structural stability after printing. The height, width, and depth of the block were determined by the maximum probability that the cubes would have no preferred orientation when dispersed in liquid. The dimensions of the capillary were chosen to maximize the uniformity of restricted signal. The main challenge of the 3D printing task was balancing the trade-off between the need for small hatching (raster) and slicing (interlayer) distances in a reasonably short printing time. For instance, the printing time per block was under 10 minutes for samples A and B in Figure 2. However, these samples had unacceptably rough inner capillary surfaces and noticeably porous inter-capillary walls. Inner capillary walls were made much smoother and without any noticeable porosity by compiling printing jobs with two added contouring lines (Sample C in Figure 2). Compared to Sample A, this also made printing time two times slower resulting in a printing time of 60 hours for a 15 mm x 15 mm array. Because MRI is sensitive to surface roughness of the samples, we chose the longer printing time to achieve a smoother cylinder wall. It was previously shown that plasma treating PMMA significantly decreases the observed water contact angle [35] making the surface more hydrophilic. This treatment significantly helped overcome the capillary pressure to load the phantom with water. However this increase in hydrophilicity may have lead to the appearance of a ‘bound’ water population, discussed more during the dPFG model analysis.

### 4.2. Usefulness for Diffusion MRI Method Validation

These arrays of aligned capillaries conform to analytical models of restricted diffusion in cylindrical pores [2, 36, 37, 38] to which NMR or MRI experiments could be compared. In these cases an average pore diameter could be estimated using a fitting procedure. If wafers were stacked having a known distribution of pore diameters or IDs, it would be possible to construct a diameter distribution phantom, such as one previously made using glass capillary arrays (GCA) [13] In this case, the diameter distribution could be estimated using an empirical distribution, although it would be useful to employ a known parametric model to determine whether experimental and model results are consistent. A limitation to this phantom preparation is that water is present within the capillaries and surrounding the blocks. Previous phantoms like the GCA found in [13] were often loaded with *H*_2_*O* and then dispersed in an *H*_2_*O* immiscible fluid or *vice versa*. A desired property to incorporate in the future is to control inter-pore exchange, for example as a means to calibrate a diffusion exchange spectroscopy (DEXSY) NMR or MRI experiment. A clever way to repurpose this phanton for a DEXSY experimental calibration is to consider the solvent domain at the ends of the capillaries in which water or other solvent in the tubes is close to the pore opening, and which has time to move into the free water compartment from the pores and *vice versa* during the mixing time of the experiment [39]. Mixing time dependence could also be studied, as well as pore hopping processes. Such experiments can be feasibly done using this phantom.

### 4.3. Low Magnetic Susceptibility

It is known that magnetic susceptibility effects can be significant in porous structures if there is a mismatch in susceptibility between the solvent and porous medium. Such is the case in GCAs. It exists here in this polymeric phantom to a much lesser extent due to the smaller susceptibility difference between plastic and water. The IP-S photo resin is primarily comprised of PMMA, a material previously shown to have magnetic susceptibility close to water, *X*_*PMMA*_*− X*_*H*2*O*_ = *−*0.036*ppm*, and was evaluated as a highly compatible MR material [40]. Classical analytical and numerical finite element method (FEM) models indicate that there would be an induced magnetic field within and around a hollow cylindrical tube which would depend on the orientation of the tube with respect to the main applied magnetic field [41, 42]. As such, capillaries that are all parallel to the main magnetic field should see no change in induced magnetic field, whereas when the tubes are aligned perpendicular to the main magnetic field, the maximum change in net magnetic field will be observed. When the aspect ratio of the tubes is large, one would expect the magnetic field within the tube to be uniform. This magnetic field dispersion is expected to cause a spread in the observed Larmor frequencies if the material had an intrinsically high *X* with respect to water. We consider the *X* of the phantom in the case in which the blocks are suspended in solution and create a randomly oriented emulsion as compared with all organized into parallel wafers or plates. An analytical solution for the orientation dependent effective frequency perturbation, ^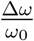^for a hollow cylinder model is presented in [43] for an infinite cylinder with an inner radius, *r*_*i*_ and an outer radius, *r*_*o*_, with the azimuthal angle *ϕ* and the altitude angle *θ* is reproduced in the 4.

The authors considered the cylinder to be comprised of the bi-lipid layer or a myelin sheath. In the case of our phantom *r*_*i*_ = 10 *µm, r*_*o*_ = 12 *µm*, or the pitch of the array. We consider solely the mobile spins because immobile spins, the spins associated with the phantom, a solid polymer material, have extremely short *T*_2_’s and do not contribute to DDE measurements and we can approximate the contribution of the region *r*_*i*_ *< r < r*_*o*_ to zero. So the contribution to an orientation-dependent frequency perturbation can only come from spins outside the capillaries. The proton spins associated with water outside the 3D printed blocks are orientationally averaged and unrestricted so we do not consider them. If we estimated the affect of neighboring hollow cylinders considering the spins near the inside wall, 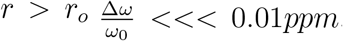. This small perturbation is unobserved on our current instrument due to the inherent resolution of the RF coil. To confirm this behavior, individually oriented blocks were investigated via PRESS, voxelwise spectroscopy, experimentally showing that the susceptibility differences between the *H*_2_*O* and the phantom material are negligible as the Larmor frequency does not vary significantly with orientation. To further show the lack of susceptibility artifacts influencing image quality, EPI data were acquired with 2,4,8 segments as shown in 4 A, B, and C respectively. When comparing the 6 and 8 segment images to the 2 segment image they have a have structural similarity (SSIM) indexes[44] of 0.9984 and 0.9661, respectively demonstrating the absence of susceptibility artifacts in that sequence, and the versatility of acquisitions that can be employed with this phantom.

### 4.4. Investigating Double Pulse Field Gradient Sequences

Our group and others have presented several diffusion methods that take advantage of DDE in order to enhance sensitivity to both the size and shape of restriction within the voxel[45]. As a first pass, we chose to apply a double-PFG filter before an EPI acquisition in order to test the method’s ability to characterize the phantom. First, let us discuss the results for the aligned phantom presenting in 7 where first gradient pulse was aligned with the capillary major axis and only the azimuth angle between the first and second gradient pulse was varied, all other experimental variables were held constant, see 3. Each voxel has a resolution of (50 × 50 × 200 *µm*), with the slice thickness chosen to ignore the top and bottom edges of the blocks. The short mixing time, *τ*_*m*_ = 0, data was fitted using the model described above. The resulting fit to the data is shown in 7, where data with widely varied parameters *G, ψ*, & *θ* were fit using mixed Gaussian and restriction model components. The model trends well with the data although the estimated diameter, 8.58 *µm*, is less the ground truth measurement from SEM. This could be attributed to the ‘channel’ like structures that are depicted in 3 that are approximately 50*µm* in width. In 30 *ms* a water molecule travels 13 *µm*, which means all of the restricted water should have hit a capillary wall at least once, however some of the unrestricted water could also have encountered the boundary of the channels. Increasing this diffusion time would eliminate the ‘free’ or Gaussian-like signal, *≈* representing 0.38 of the total fraction of attenuating signal. For the orientationally-distributed phantom a dPFG experiment was performed with 3 shells, incremented angles, and gradient strengths at a 3D voxel resolution of 100 *µm* cubed. We decreased the DTI resolution used for this experiment from 50 *µm* to 100 *µm* voxels, the apparent FA, (8, A), is underestimated, resulting from “powder averaging”, a known issue for DTI, thus the voxels used for the diameter estimate had FAs *>* 0.45, (8, B) although the SI FA high-resolution image for the same phantom shows the block with FA closer to 0.8, as expected. The resulting estimated diameter map in, (8, C), shows that the mixed Gaussian model (with one restricted and one free population) does not necessarily capture complexity of the phantom. A bound population may exist in the phantom due to the hydrophilicity of phantom’s microcapillary walls. For long mixing time the orientationally-averaged signal for water diffusing with an apparent diffusivity, *D*_0_, in an infinite cylinder with radius, *a* was presented in [46, 47] by exploiting separation of variables. This can be represented by *E*(*g, ϕ*), see below

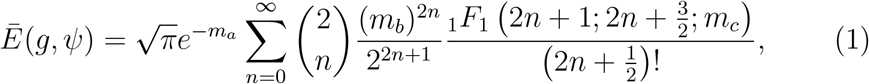

where *m*_*a*_,*m*_*b*_ are dependent on the experimental parameters Δ, *δ,ψ* and the estimated parameters *D*_0_, *r*_*c*_*ylinder*, described fully in Appendix A.

Using this model the angle dependent results were fit yielding the following parameters displayed in 5. Initially no bound water population was included in the fit yielding a low estimated diameter, 7.80 *µm±* 0.08. Forthe both the free/bound/restriction and free/restriction models the apparent diffusivity was set to 1.85 *m*2*s*^−1^ based on the known free diffusivity of water at the sample temperature, while the fraction of freely diffusing spins, capillary diameter, and signal intensity were optimized using least squared analysis. The model without a bound population severely underestimated the expected diameter while inclusion of the bound water produces a diameter closer to microscopic ground truth. We suspect that the plasma treatment of the blocks before loading with water caused a layer of bound water to the surface of the blocks with increased hydrophilicity. To investigate this magnetization transfer FLASH3D experiments were performed at 50 micron resolution, (see SI), however as shown by the PRESS spectra there is not a broad hump of immobile water convolved with a sharp Lorentzian, as is the case for most tissue spectra, and the MT images did not result in a significant difference from off-resonance excitation or indication of a bound population. However, *T*_2_ weighted map **??**, unexpectedly shows that the water surrounding the blocks has a much shorter *T*_2_ than the water inside the block. This could be due to a hydrophilic layer on the outside surface of the phantom. Bound water does not need to participate in a hydrogen bond, however the interaction with the surrounding media needs to be ‘strong’ enough to distinguish its mobility from the bulk water behavior[48].

While the randomly oriented phantom can exhibit both macroscopic isotropy and microscopic anisotropy, the SEM microscopy images reveal the ground truth to be 10 *µm* while our short and long *τ*_*m*_ dPFG fits to the MRI experiments underestimate the capillary diameter. We considered first that our phantom may contain a surface bound water population (see T1 images in SI). A significant amount of the signal decay is due to freely diffusing water, for the model explored in this paper *D*_0_ and *d* are highly correlated due to this. To weaken this correlation we plan to change the liquid filling the phantom to a higher viscosity inert small molecule or sol-gelling material that can still be loaded into the small capillaries, and use phantom designs that reduce the free water contribution.

## 5. Conclusion

High-resolution 3D printing is opening up new opportunities for phantom fabrication including the creation of reproducible, durable, ground-truth structural phantoms that can mimic complex structural features of tissue. Here, we have shown that ground-truth phantoms with microscopic pore sizes can be fabricated via 3D printing and filled with water resulting in a microscopically anisotropic phantom that does not suffer from susceptibility artifacts. Although a single pore size was presented here, this method can also produce controlled distributions of pores sizes, varied wall permeabilities, and controlled orientation distributions, which are needed to improve and validate methods that seek to characterize these features in tissues. In the case of diffusion exchange, which is increasingly being understood as a process occurring in gray matter [49] we plan to use this phantom to vet diffusion exchange spectroscopic (DEXSY) [50] methods as in [39]

## 6. Acknowledgments

A portion of this research was conducted at the Center for Nanophase Materials Sciences, which is a DOE Office of Science User Facility. The authors thank Drs Ye Sun and Christopher Bleck for their electron microscopy technical expertise. SEM imaging was performed on a Hitachi instrument maintained by the NIH, National Heart, Lung, and Blood Institute (NHLBI). VJW and PJB were supported by the Intramural Research Program (IRP) of the *Eunice Kennedy Shriver* National Institute of Child Health and Human Development (NICHD). MK and DB were supported by the Center for Neuroscience and Regenerative Medicine (CNRM) under the auspices of the Henry M. Jackson Foundation (HJF). DB was also supported in part by the IRP of the National Institute on Aging. Fabrication of the MRI phantoms were conducted as part of a user project at the Center for Nanophase Materials Sciences (CNMS), which is a US Department of Energy, Office of Science User Facility, Oak Ridge National Laboratory.

## Appendix

### A. Appendix

Here, we provide the derivation of the double-PFG MR signal intensity for an isotropic distribution of infinite cylinders. We stress that our results can be used even in the presence of orientational coherence [51] by employing an adequate orientational-averaging scheme [52] on multi-directional double diffusion encoding acquisitions. Herberthson et al. [53] derived the orientationally-averaged MR signal for general gradient waveforms under the assumption of free diffusion within each compartment. These results were extended by Yolcu et al., [47] who provided explicit expressions for singleand double diffusion encoding measurements performed on a collection of confinement tensors [54, 55]. The exact expression for the signal attenuation for one confinement tensor is given by [47]

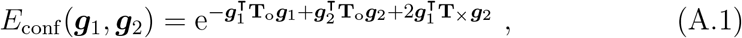

where ***g***_1_ and ***g***_2_ denote the product of the gyromagnetic ratio and the gradient vectors in the first and second blocks, respectively. We would like to write down the same expression for cylinders, i.e., find the corresponding tensors **T**_o_ and **T**_×_ for infinite cylinders of radius *a* and bulk diffusivity *D*_0_.

Consider a single such cylinder oriented along the unit vector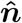. The ***g***_*i*_, (*i* = 1, 2) vectors can be decomposed into directions parallel and perpendicular to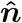, as in Liat Avram et al. [**?**] given by 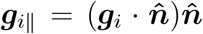 and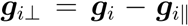. Due to separation of variables, the signal attenuation for diffusion within the cylinder can be written as

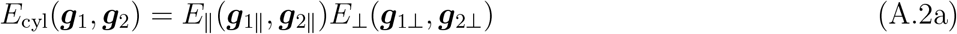

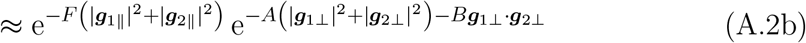

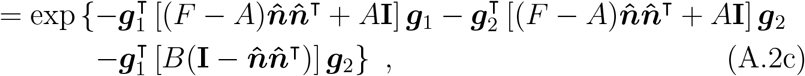

where **I** denotes the 3 × 3 identity matrix, and the expressions for *F, A*, and *B* are provided by Ozarslan and Basser [51] and reproduced below for completeness:

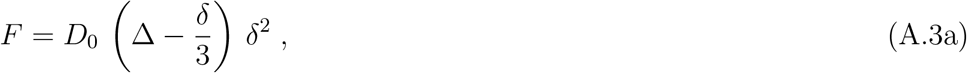

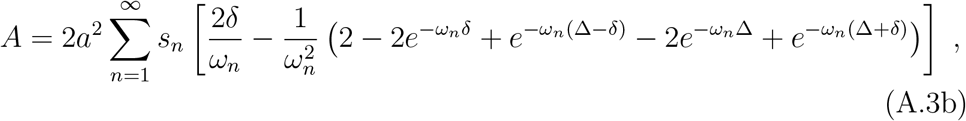

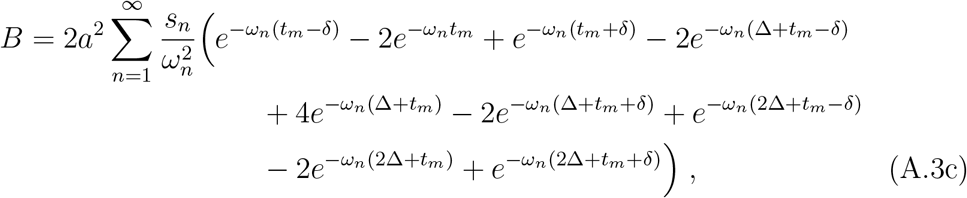

where

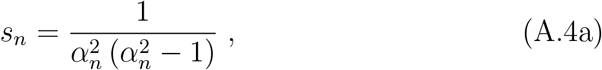

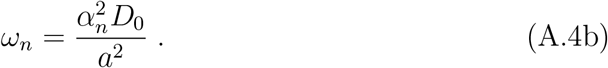

Here, *α*_*n*_ is the *n*th zero of the derivative of the first order Bessel function, i.e., the expression 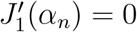 is satisfied.

Note that the expression for the signal from one cylinder, Eq. (A.2c), is indeed of the form in (A.1) with the substitutions

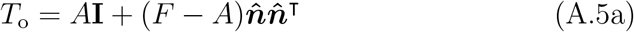

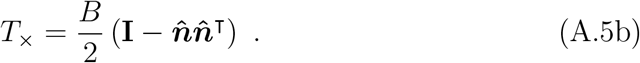

Thus, the signal for an isotropic distribution of a cylinder can be obtained by adapting Eq. (25) in the work of Yolcu et al. [47] to our problem, yielding the expression

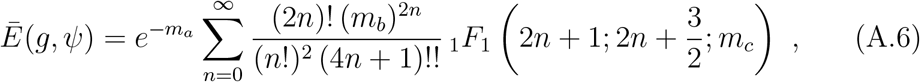

where *g* = _|_ ***g***_1_ = ***g***_2|_ is the gyromagnetic ratio multiplied by the gradient strength, which is the same in both blocks of the double-PFG sequence.

The angle between the gradient directions is denoted by *ψ* while _1_*F*_1_ is the confluent hypergeometric function. Moreover,

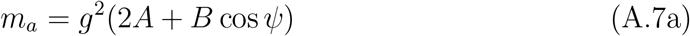

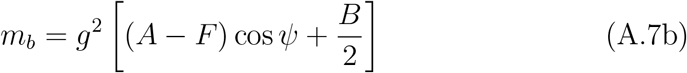

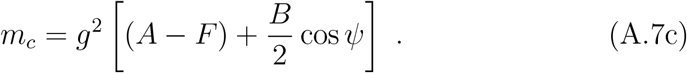

