## SUPPLEMENTARYINFO for "Novel Pore Size-Controlled, Susceptibility Matched, 3D-Printed MRI Phantoms"

### 1. Supporting Figures

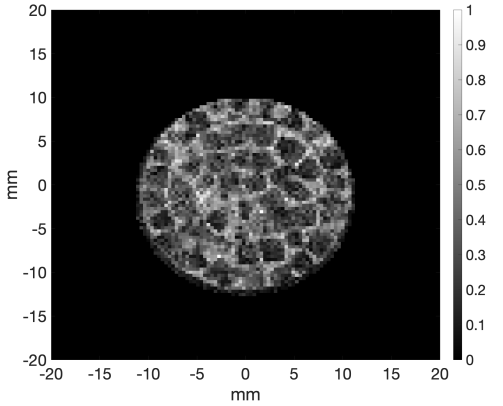


Figure 1: Inversion recovery weighted FLASH3D map with Inversion time = 13 ms

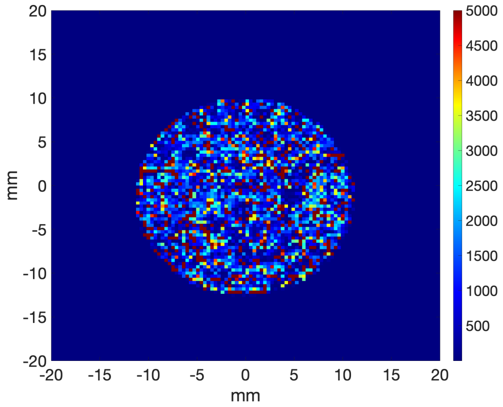


Figure 2: Inversion recovery weighted FLASH3D map for slice 25.

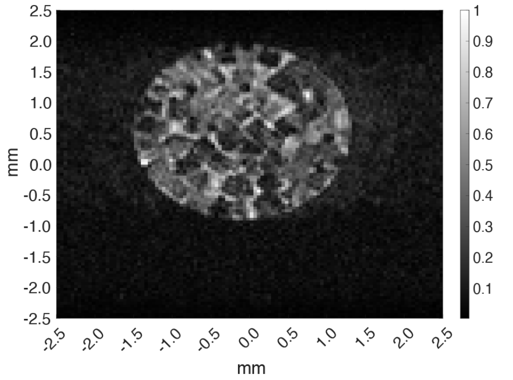


Figure 3: Diffusion Filtered (b-value 300) T1 map using a DWI-Inversion Recovery time of 200 ms with EPI Acquisition for slice 25

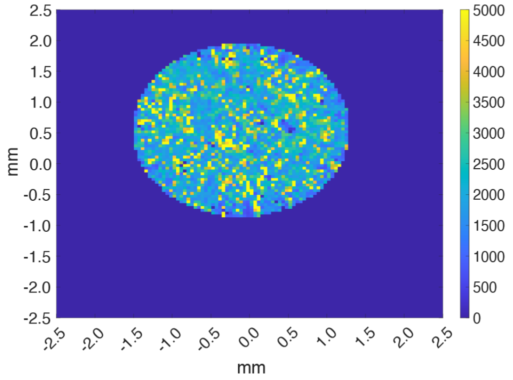


Figure 4: Diffusion Filtered (b-value 300) T1 map using a DWI-Inversion Recovery EPI Acquisition for slice 25

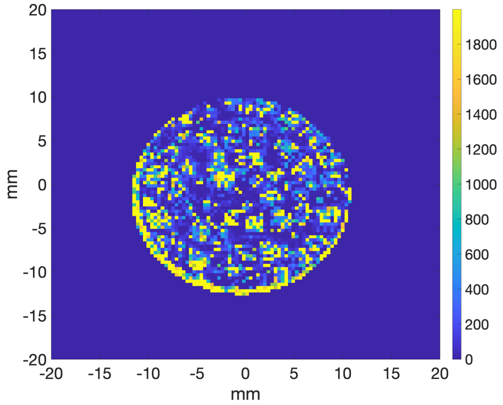


Figure 5: Axial slice 20 of a FLASH3DMAP with multi gradient echo acquisition where a custom MATLAB code was used to calculate the $T_2*$ MAP.
